# Robust neutralization assay based on SARS-CoV-2 S-bearing vesicular stomatitis virus (VSV) pseudovirus and ACE2-overexpressed BHK21 cells

**DOI:** 10.1101/2020.04.08.026948

**Authors:** Hua-Long Xiong, Yang-Tao Wu, Jia-Li Cao, Ren Yang, Jian Ma, Xiao-Yang Qiao, Xiang-Yang Yao, Bao-Hui Zhang, Ya-Li Zhang, Wang-Heng Hou, Yang-Shi, Jing-Jing Xu, Liang-Zhang, Shao-Juan Wang, Bao-Rong Fu, Ting Yang, Sheng-Xiang Ge, Jun Zhang, Quan Yuan, Bao-Ying Huang, Zhi-Yong Li, Tian-Ying Zhang, Ning-Shao Xia

## Abstract

The global pandemic of Coronavirus disease 2019 (COVID-19) is a disaster for human society. A convenient and reliable *in vitro* neutralization assay is very important for the development of neutralizing antibodies, vaccines and other inhibitors. In this study, G protein-deficient vesicular stomatitis virus (VSVdG) bearing full-length and truncated spike (S) protein of SARS-CoV-2 were evaluated. The virus packaging efficiency of VSV-SARS-CoV-2-Sdel18 (S with C-terminal 18 amino acid truncation) is much higher than VSV-SARS-CoV-2-S. A neutralization assay for antibody screening and serum neutralizing titer quantification was established based on VSV-SARS-CoV-2-Sdel18 pseudovirus and human angiotensin-converting enzyme 2 (ACE2) overexpressed BHK21 cell (BHK21-hACE2). The experimental results can be obtained by automatically counting EGFP positive cell number at 12 hours after infection, making the assay convenient and high-throughput. The serum neutralizing titer of COVID-19 convalescent patients measured by VSV-SARS-CoV-2-Sdel18 pseudovirus assay has a good correlation with live SARS-CoV-2 assay. Seven neutralizing monoclonal antibodies targeting receptor binding domain (RBD) of SARS-CoV-2-S were obtained. This efficient and reliable pseudovirus assay model could facilitate the development of new drugs and vaccines.

## Introduction

As of April 5, 2020, over 1.2 million cases of coronavirus disease 2019 (COVID-19) have been reported, including 68,000 deaths. The spread of the SARS-CoV-2 virus is difficult to control due to the highly contagious and asymptomatic infection. There is an urgent need to develop vaccines or therapeutics against SARS-CoV-2 infection. However, SARS-CoV-2 culture and assays need to be carried out in biosafety level-3 laboratory, which limits the efficiency of research and development (R & D). Widely available and convenient functional assays, such as cell-based assays for neutralizing quantification, are essential for vaccine and therapeutic antibody R & D ^[1–3]^.

The SARS-CoV-2 belonging to the genus of betacoronavirus with envelope ^[4]^. The glycosylated spike (S) protein is the major surface protein and is responsible for receptor binding and fusion of the viral membrane with cellular membranes^[5, 6]^. Studies has showed that SARS-CoV-2 S and SARS-CoV S bind with the same receptor, human angiotensin-converting enzyme 2 (hACE2), with similar affinities^[7–9]^. Therefore, the studies of SARS-CoV neutralization assays, which based on pseudovirus bearing the spike protein of SARS-CoV, would provide valuable experience and reference for the SARS-CoV-2 assays. In the past years, pseudotyping system based on vesicular stomatitis virus (VSV) was reported to produce pseudotypes incorporating the envelope protein of heterologous risk group-3 (RG-3) or RG-4 viruses, such as Ebola virus, SARS coronavirus, MERS coronavirus et al., in which the VSV G gene is deleted (VSVdG) and the gene encoding GFP, luciferase or other reporter genes was integrated ^[10–12]^. Pseudotyped viruses provide a safe viral entry model because of their inability to produce infectious progeny virus. VSV assembly occurs at the plasma membrane and involves budding of virions from the cell surface. During budding, VSV acquires an envelope consisting of a lipid bilayer derived from the plasma membrane and spike proteins consisting of trimers of the VSV glycoprotein (VSV-G). When the VSV-G is absence and the glycoprotein from heterologous virus is complacently expressed in cells infected with rVSV-dG, the glycoprotein of heterologous virus could be assembled into the VSV membrane.

Recently, Michael Letko et al. used VSVdG-luc bearing SARS-spike chimeras to study the cell entry and receptor usage for SARS-CoV-2 and other lineage B betacoronaviruses^[13]^. Jianhui Nie et al. have successfully constructed a pseudovirus neutralization assay for SARS-CoV-2, which consist of pseudotyped VSV bearing the full-length spike protein of SARS-CoV-2 and Huh7 cell^[12]^. Lentiviral pseudotype bearing the truncated spike protein of SARS-CoV-2 was also constructed and used to study the virus entry and its immune cross-reactivity with SARS-CoV ^[6]^. Noteworthy, the correlation between pseudovirus system and live virus system hasn’t been verified. And the virus package and infection efficiency is also one of the limiting factors for high-throughput in vitro neutralization assay.

It was reported that a pseudotyped VSV and retrovirus bearing a SARS-CoV-S protein variant with a truncation in the cytoplasmic tail had much higher infection efficiency than that with full-length S protein^[11, 14]^. In this study, the VSVdG pseudotyped with full-length SARS-CoV-2-S protein or truncated SARS-CoV-2-Sdel18 protein with C-terminal 18 aa truncation were compared, we found that the infection efficiency of VSV-SARS-CoV-2-Sdel18 was much higher than VSV-SARS-CoV-2-S. Among the cell lines used in this study, BHK21-hACE2 stably expressing human ACE2 was most sensitive for infection, while Vero-E6 was most suitable for virus package. A neutralization assay for antibody screening and validation was established based on VSV-SARS-CoV-2-Sdel18 and BHK21-hACE2 cell, and 7 strains of neutralizing monoclonal antibodies targeting receptor binding domain (RBD) of SARS-CoV-2-S were obtained. Most importantly, the correlation between pseudovirus system and live virus system was verified.

## Materials and Methods

### Cells and Samples

Vero-E6 (American Type Culture Collection [ATCC], CRL-1586), Vero (ATCC, CCL-81), BHK21 (ATCC, CCL-10) and 293T (kindly gifted by Dr. Jiahuai Han) cells were maintained in high glucose DMEM (SIGMA-ALDRICH) supplemented with 10% FBS (GIBCO), penicillin (100 IU/mL), streptomycin (100 μg/mL) in a 5% CO2 environment at 37°C and passaged every 2 days. BHK21-hACE2 cell was developed by stably transfection of hACE2-expressing plasmid following puromycin resistance selection.

SARS-CoV-2 spike specific antibody was generated by immunized Balb/c mouse with SARS-CoV-2 spike protein (RBD, Fc Tag, 40592-V05H, Sino Biological inc). MAbs against SARS-CoV-2 RBD were produced using hybridoma technology and were characterized by pseudotyped virus assay.

Eighteen SARS-CoV-2 serum samples from convalescent patients were generously provided by Mr. Zhi-Yong Li from The First Hospital of Xiamen University. Written informed consents were obtained from all the volunteers.

### Plasmids and pseudovirus

To construct VSV pseudovirus carrying the spike protein of SARS-CoV-2, the spike gene of Wuhan-Hu-1 strain (GenBank: MN908947) was codon-optimized for expression in human cells and cloned into the eukaryotic expression plasmid pCAG to generate recombinant plasmid pCAG-nCoVS. The spike gene mutant of SARS-CoV-2 that with C-terminal 18 aa truncation was also cloned into plasmid pCAG and generate pCAG-nCoVSde18. Plasmid pCAG-nCoVS and pCAG-nCoVSde18 were transfected into Vero-E6 respectively. 48 hours post transfection, the VSVdG-EGFP-G (Addgene, 31842)^[15]^ virus were inoculated to the cell expressing SARS-CoV-2 spike protein or SARS-CoV-2 Sde18 truncated protein and incubated for one hour. Then removed VSVdG-EGFP-G virus in supernatant and added anti-VSV-G rat serum to block the infectivity of residual rVSVdG-G. The progeny virus would be enveloped by SARS-CoV-2 spike protein or SARS-CoV-2 Sde18 truncated protein to generate VSV pseudovirus carrying different spike protein of SARS-CoV-2, as shown in Figure 1. The supernatant was harvested at 24 hours post rVSVdG-G infection and then centrifuged and filtered (0.45-μm pore size, Millipore, SLHP033RB) to remove cell debris and stored at −80°C until use. The virus titer was determined by counting the number of GFP positive cells after infected BHK21-hACE2 with gradient diluted supernatant.

**Figure 1:**
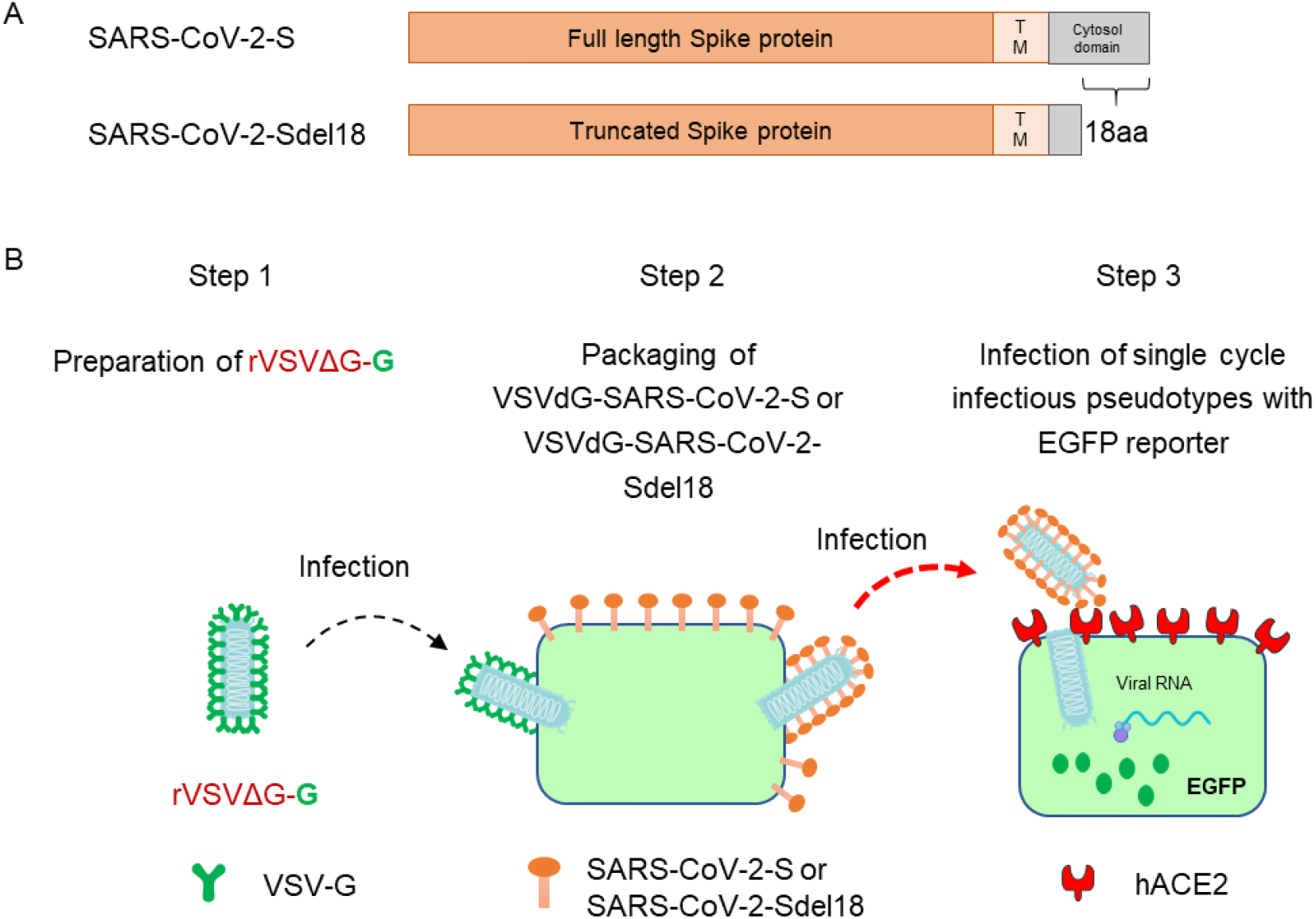
Generation of VSV pseudotypes bearing SARS-CoV-2 spike proteins. (A) The difference between SARS-CoV-2-S and SARS-CoV-2-Sde18. (B) The procedure of producing VSV pseudotypes bearing SARS-CoV-2 spike proteins.

### Pseudotype-based neutralization assay

BHK21-hACE2 cell was seeded previously. Culture supernatant of monoclonal hybridoma cells, gradient diluted purified antibodies or serum of convalescent patient were mixed with diluted VSV-SARS-CoV-2-Sdel18 virus (MOI=0.05) and incubated at 37°C for 1h. All the sample and virus were diluted using 10%FBS-DMEM. The mixture was added to seeded BHK21-hACE2 cell. After 12h incubation, fluorescence images were obtained by using Opera Phenix or Operetta CLS equipment (PerkinElmer). For quantitative determination, fluorescence images were analyzed by Columbus system (PerkinElmer) and the numbers of GFP-activated cell for each well were counted to represent infection performance. The reduction (%) of mAb treatment GFP-activated cell numbers in comparing with non-treated control well was calculated to show the neutralizing potency.

### Live SARS-CoV-2-based neutralization assay

Seeded Vero cells in 96-well plates. The next day, changed the 10%FBS-DMEM medium to 2%FBS-DMEM and transferred the cell to P3 laboratory for subsequent analysis. The serum of convalescent patient and Live SARS-CoV-2 (C-Tan-nCoV Wuhan strain 01) were diluted using 2%FBS-DMEM. The mixture of gradient diluted serum and virus were incubated at 37°C for 1h and added to seeded Vero cell. After incubating at 37°C for 48h, observed the CPE and took 100μL of the culture supernatant for nucleic acid extraction. Took 5μL of nucleic acid to perform real-time fluorescence RT-PCR reaction on ABI Q5 fluorescence quantitative PCR instrument. Calculated the virus TCID50 in the sample according to CT value of the sample and standard curve. Compared the TCID50 of serum neutralized sample to control and calculated the inhibition rate.

### Statistic

The inhibition rates of mAb and serum were calculated according to the decrease of GFP positive cell number (for pseudotype-based neutralization assay) or decrease of RNA levels (for live SARS-CoV-2-based neutralization assay). The IC50 (the half maximal inhibitory concentration) or ID 50 (plasma/serum dilutions causing a 50% reduction of GFP-positive cell number or RNA level) values were calculated with non-linear regression, i.e. log (inhibitor) vs. normalized response – Variable slope, using GraphPad Prism 7.00 (GraphPad Software, Inc., San Diego, CA, USA). The correlation between analysis result of pseudovirus system and live virus system was analysed by linear regression using GraphPad Prism 7.00.

## Results

### The combination of VSV-SARS-CoV-2-Sdel18 and BHK21-hACE2 cells showed higher efficiency

To compare the infection efficiency of pseudotypes bearing different glycosylated protein, single cycle infectious recombinant VSVdG-EGFP-G was prepared and used to infect Vero-E6 cells transiently expressing full-length SARS-CoV-2-S protein and truncated SARS-CoV-2-Sdel18 protein that with C-terminal 18 aa truncation respectively, pseudotyped VSV-SARS-CoV-2-S and VSV-SARS-CoV-2-Sdel18 was obtained following the procedures described in methods (Figure 1). Vero-E6, BHK21, BHK21-hACE2 and 293T cells were seeded to compare the infection efficiency of pseudotypes. The infection efficiency of VSV pseudotypes bearing truncated Sdel18 was much higher than that with full-length S protein. Similar results were observed in BHK21-hACE2 and Vero-E6 cells, among which the BHK21-hACE2 stably overexpressing hACE2 was most sensitive, whereas the BHK21 cells and 293T cells were almost resistant to infection (Figure 2). The number of GFP-positive cells in VSV-SARS-CoV-2-Sdel18 infected BHK21-hACE2 was about 70 times more than that in VSV-SARS-CoV-2-S infection group, and 1.26 times lower than that in the VSVdG-G infection group (Figure 2). The BHK21-hACE2 infected with VSV-SARS-CoV-2-Sdel18 also formed more symplasm than cell infected with VSV-SARS-CoV-2-S (Supplementary figure 1). The results suggested that the combination of VSV-SARS-CoV-2-Sdel18 and BHK21-hACE2 cells could be a better alternation for establishing pseudotyped SARS-CoV-2 entry and neutralization assays.

**Figure 2:**
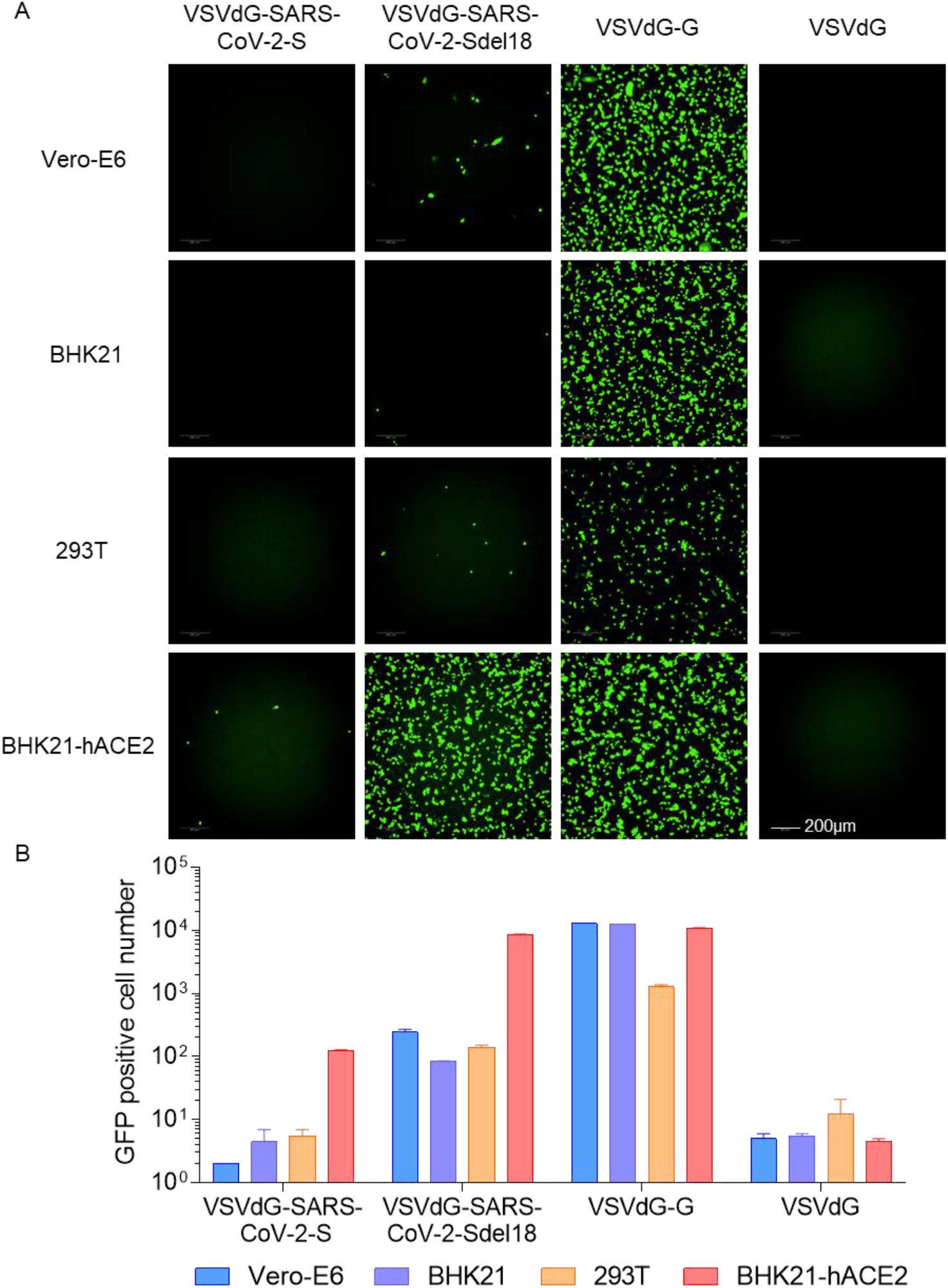
Comparison of the infectious efficiency of pseudotypes in various cell lines. VSVdG virus bearing spike protein of SARS-CoV-2 or G protein of VSV were harvested and the infectivity of these recombinant virus were tested in different cell lines, including Vero-E6, BHK21, 293T and BHK21-hACE2. The fluorescence was detected (A) and GFP positive cell number (B) was counted using Opera Phenix 12 h post infection.

### Vero-E6 cells packaged the VSVdG-SARS-CoV-2-Sdel18 with highest efficiency

Viral packaging efficiency is one of the limiting factors for high-throughput in vitro neutralization assay. To select a cell line that is most appropriate for pseudotypes production, we compared the viral titers of VSVdG-SARS-CoV-2-Sdel18 packaged by Vero-E6, BHK21 and 293T cells. The results showed that the Vero-E6 formed more syncytia than the other two cell lines during the packaging, and also produced highest viral titer (Figure 3), which was about 3×10 ^5^ infectious particles per miniliter.

**Figure 3:**
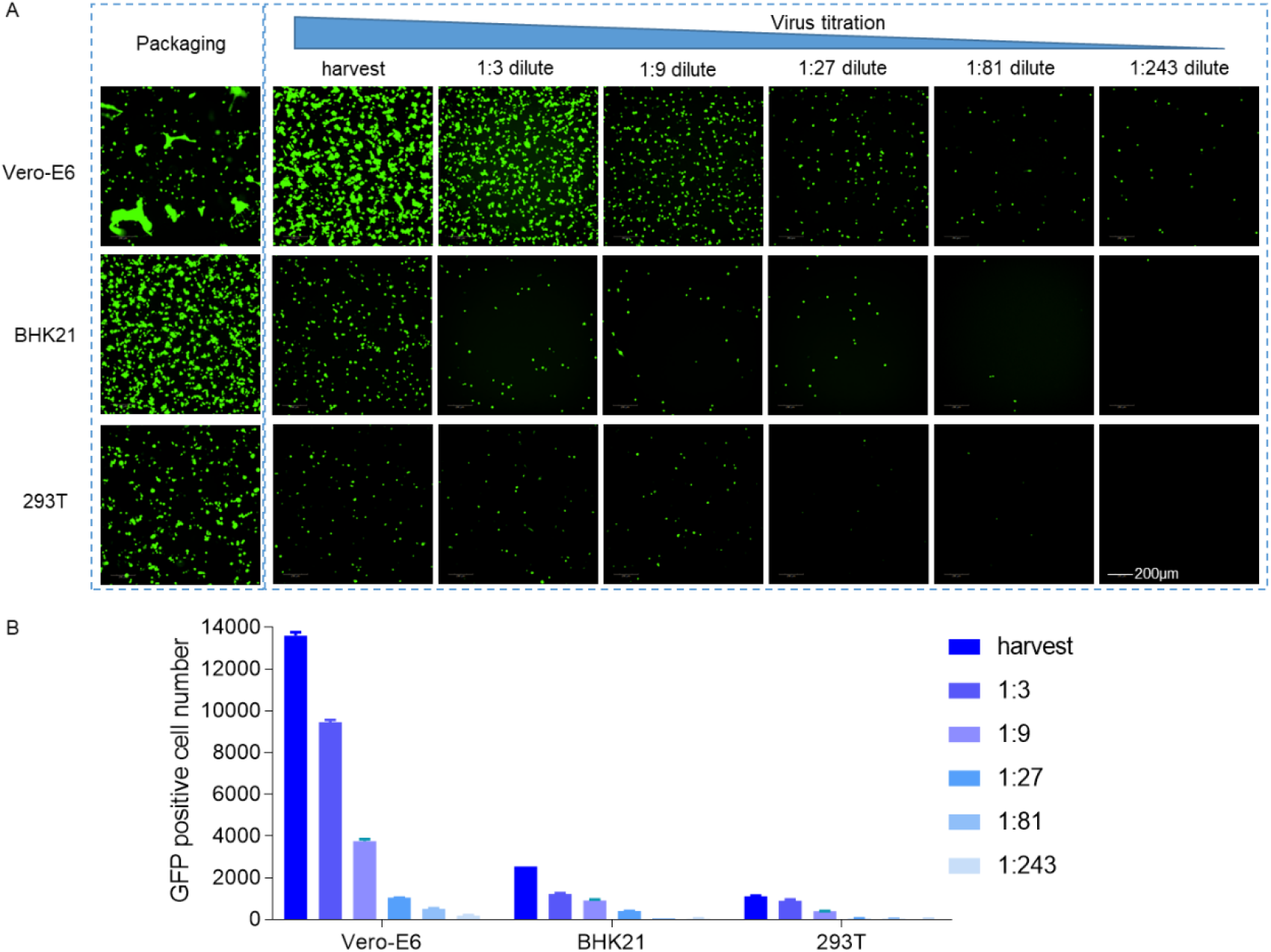
Comparison of the packaging efficiency of VSVdG-SARS-CoV-2-Sdel18 in various cell lines. Vero-E6, BHK21 and 293T cells were used to package VSVdG-SARS-CoV-2-Sdel18 virus. The left picture shows the cell used to package recombinant virus, recorded 48h post infection of VSVdG-EGFP-G. The right figures show the infectivity of virus produced by three cell lines. The harvested virus was diluted and tested in BHK21-hACE2 cell. The fluorescence was detected (A) and GFP positive cell number (B) was counted using Opera Phenix 12 h post infection.

### Time course of EGFP expression after VSVdG-SARS-CoV-2-Sdel18 infection

To determine the optimal incubation period for detection of VSVdG-SARS-CoV-2-Sdel18 infection, a time-course monitoring of EGFP expression was performed using Opera Phenix High Content Screening System. EGFP expression could be detected as early as 8 hours post infection, and the number of GFP-positive cells gradually increased, reaching the plateau phase within 12 hours, and remained stable between 12-24 hours (Figure 4). Whereas the fluorescence intensity was increased continuously. As pseudoviruses are unable to produce progeny viruses, the infected cell number would be a better indicator for infection efficiency. Therefore, the infection efficiency could be quantitative evaluated at 12-24 hours by counting the number of GFP-positive cells.

**Figure 4:**
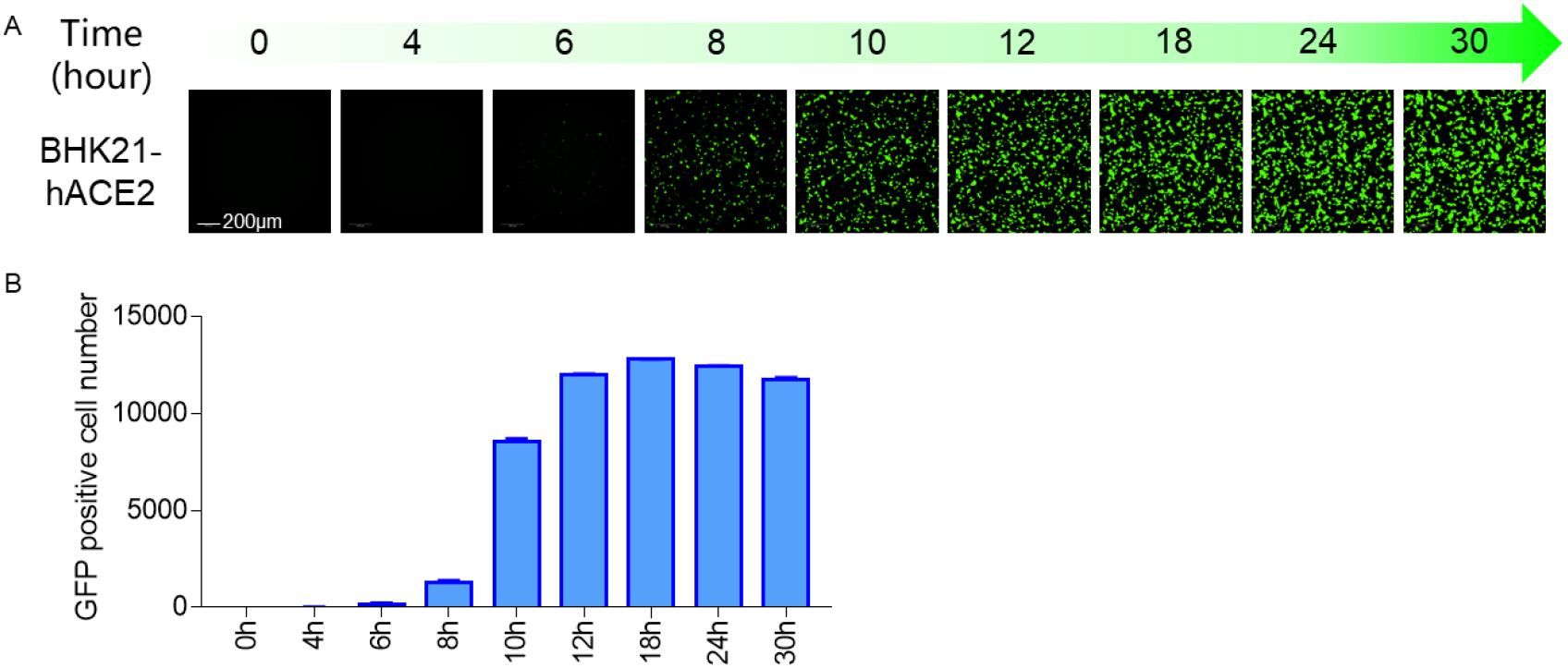
Time course of EGFP expression after VSVdG-SARS-CoV-2-Sdel18 infection. BHK21-hACE2 cell was infected with VSVdG-SARS-CoV-2-Sdel18 virus. The fluorescence was detected (A) and GFP positive cell number (B) was counted using Opera Phenix at different time point post infection.

### VSVdG-SARS-CoV-2-Sdel18 based system for screening of neutralizing mAbs

We firstly tested the response of VSVdG-SARS-CoV-2-Sdel18 based system to the treatment of chloroquine. BHK21-hACE2 was incubated with VSVdG-SARS-CoV-2-Sdel18 virus (MOI=0.05) and different concentration of chloroquine. The result indicated that the IC50 of chloroquine for VSVdG-SARS-CoV-2-Sdel18 (17.2±1.4 μM) is close to live SARS-CoV virus (8.8 ± 1.2 μM) (supplementary figure 2) ^[16]^. It also means this pseudoviruse system can mimic the entry of live virus. This system can be used for high-throughput screening of neutralizing antibodies. As shown in Figure 5A and 5B, 35 strains of mouse monoclonal antibody hybridoma culture supernatants were evaluated using the pseudovirus system, and 7 of them showed significant neutralization. The IC50 of 7 selected neutralizing antibodies for antiviral activity were also analyzed (Figure 5C) using this pseudovirus system.

**Figure 5:**
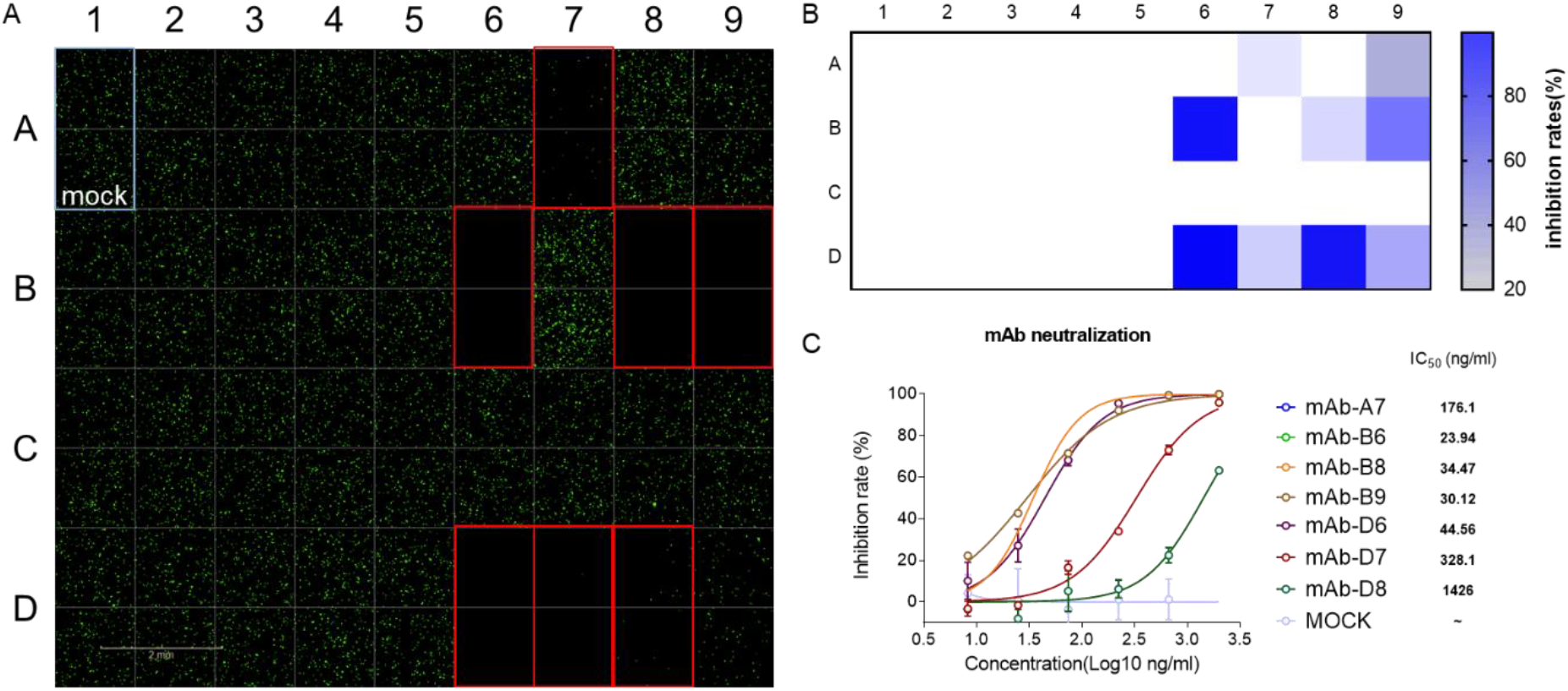
VSVdG-SARS-CoV-2-Sdel18 based system for screening of neutralizing mAbs. A Neutralizing antibodies were screened from 35 antibodies. The culture supernatant of 35 monoclonal hybridoma cells were incubated with VSVdG-SARS-CoV-2-Sdel18 virus and then added the mixture to BHK21-hACE2 cell. The fluorescence was detected using Opera Phenix 12 h post infection. The GFP positive cell number was also counted to calculate the inhibition rate (B). C The IC50 of 7 selected neutralizing antibodies for antiviral activity were also analyzed. The 7 selected neutralizing antibodies were purified and diluted to different concentration, then incubated with VSVdG-SARS-CoV-2-Sdel18 virus for an hour and added to BHK21-hACE2 cell. The GFP positive cell number was counted using Opera Phenix 12 h post infection to calculate the inhibit ratio. The IC50 was analysed by nonlinear regression.

### The correlation between VSVdG-SARS-CoV-2-Sdel18 based system and live SARS-CoV-2 system

VSVdG-SARS-CoV-2-Sdel18 based system was used to analyze the neutralizing antibodies in the serum of 18 convalescent patients. The serum was diluted 60-fold as the first concentration, and then prepared doubling dilutions (60, 120, 240, etc.). BHK21-hACE2 cell was seeded and incubated with the mixture of gradient diluted serum and VSVdG-SARS-CoV-2-Sdel18 virus (MOI=0.05). The GFP positive cell number was counted to analysed the neutralizing activity of serum (Figure 6A). Among the 18 tested sera, 5 of them show high neutralizing activity (ID_50_>1000). Similar assay was also performed using live SARS-CoV-2 system. 7 sera show high neutralizing activity (ID_50_>1000) according to live virus assay (Figure 6B). The analysis result of the pseudovirus system correlates well with live virus system (R^2^=0.639, P<0.0001, Figure 6C).

**Figure 6:**
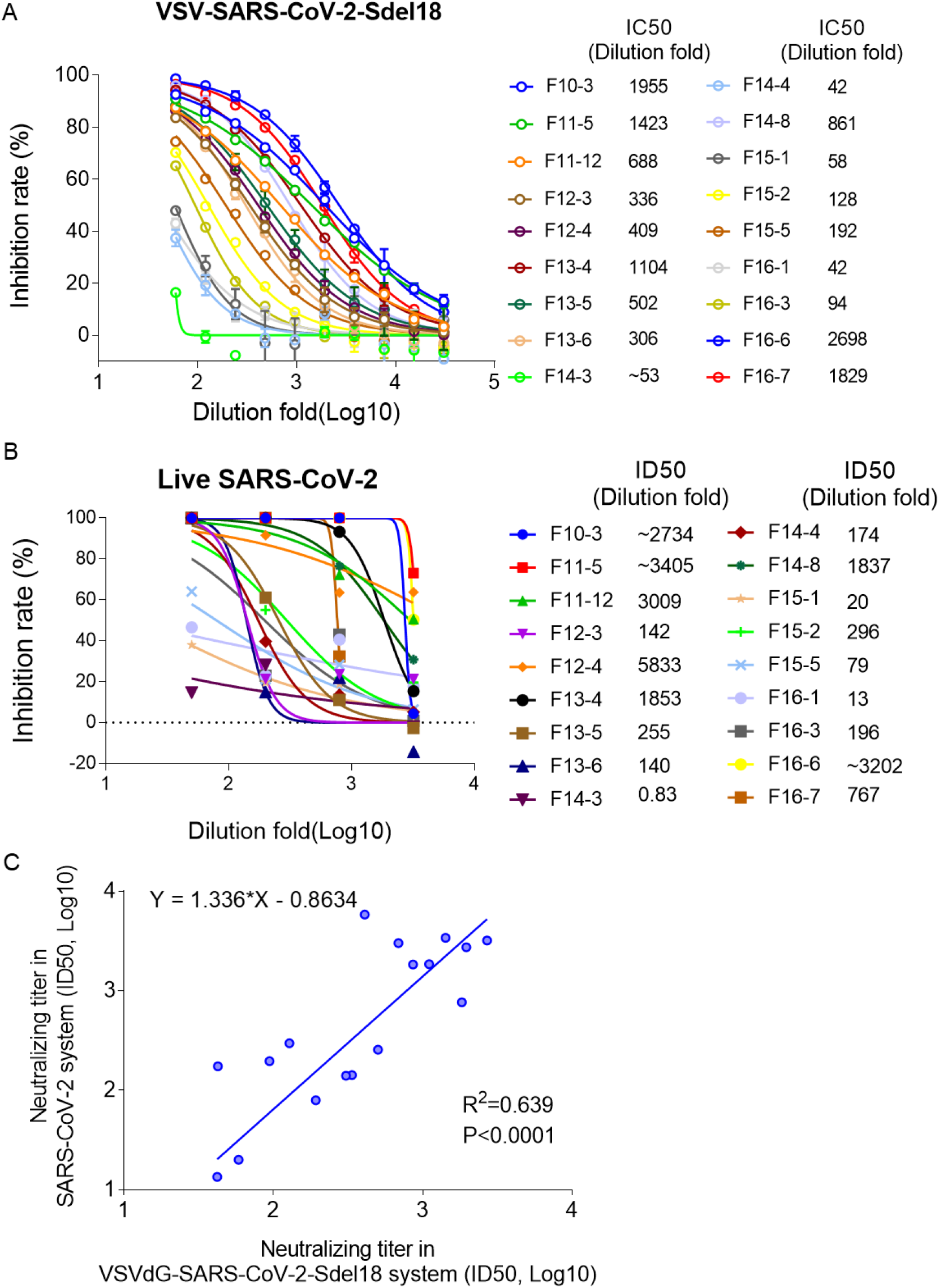
Verify the correlation between VSVdG-SARS-CoV-2-Sdel18 based system and live SARS-CoV-2 system. A Analyze the neutralizing antibodies in the serum of 18 convalescent patients using VSVdG-SARS-CoV-2-Sdel18 based system. ID50 was used to indicate the neutralizing activity of sera. B Analyze the neutralizing antibodies in the serum of 18 convalescent patients using live SARS-CoV-2 system. ID50 was used to indicate the neutralizing activity of sera. C The correlation between VSVdG-SARS-CoV-2-Sdel18 based system and live SARS-CoV-2 system.

## Discussion

The COVID-19 epidemic is spreading rapidly around the world, there is an urgent need for vaccines or therapeutic drugs. The isolation and culture of SARS-CoV-2 have to be handled in biosafety level 3 (BSL-3) facilities, which limits the research and development of drugs. A safer and more efficient assay for drugs evaluation would accelerate the development of drugs.

The pseudovirus system provides a convenient tool for the research of RG-3 viruses, it makes the evaluation of inhibitors for viral entry, neutralizing antibodies and immunized-serum much easier. Retrovirus and VSV are both common used pseudovirus system, but the pseudotype virus titer obtained with VSVdG system is generally higher than that of the retrovirus system ^[10]^. Furthermore, infection of target cells with pseudotyped VSV can be detected as early as 4 hours post infection. In this research, pseudotyped VSV carrying the spike protein of SARS-CoV-2 and reporter gene can be detected 8 h post infection through green fluorescence. It takes much shorter time than live SARS-CoV-2 virus assay. The detection of pseudotyped VSV through fluorescence is easy to operate and guarantees the high throughput of assay.

Jianhui Nie et al. have also constructed pseudovirus-based neutralizing assays, which consist of pseudotyped VSV bearing the full-length spike protein of SARS-CoV-2 and Huh7 cell, the reporter gene firefly luciferase was used to indicate the infection of pseudotyped virus ^[12]^. Noteworthy, the truncation of the cytoplasmic tail may increase fusion activity, which is consistent with the study of SIV-MuLV chimeric envelope protein^[17]^. Enhanced syncytia arising from the fusion may contribute to the improved packaging efficiency. The pseudotyped VSV with C-terminal 18 aa truncation of SARS-CoV-2 spike protein led to more obvious symplasm, comparing to pseudotyped VSV with full length SARS-CoV-2 spike protein. The truncation of spike protein also accompanied higher infectivity of pseudotyped VSV in this research.

The infection model consisting of BHK21-hACE2 cell and VSVdG-SARS-CoV-2-Sdel18 can mimic the entry of live SARS-CoV-2. We have verified the correlation between the pseudovirus assay and the live virus assay. And it is safer and time saving compared to live SARS-CoV-2. This pseudotype infection model could be applied to screening of compounds that could inhibit infection of SARS-CoV-2 and neutralizing antibody. It can also be used to evaluate the production of neutralizing antibodies after vaccine immunization. Generally, this pseudotype infection model could greatly benefit the development of new drugs and vaccines.

## Author Contribution

Hua-Long Xiong, Bao-Ying Huang, Zhi-Yong Li, Tian-Ying Zhang, and Ning-Shao Xia had full access to all of the data in the study and take responsibility for the integrity of the data and the accuracy of the data analysis. Study concept and design: Tian-Ying Zhang, Quan Yuan and Ning-Shao Xia. Acquisition of data: Hua-Long Xiong, Yang-Tao Wu, Jia-Li Cao, Jian Ma, Xiao-Yang Qiao, Bao-Hui Zhang, Yang-Shi, Jing-Jing Xu, Liang-Zhang. Ren Yang and Bao-Ying Huang performed the live virus assay. Analysis and interpretation of data: Hua-Long Xiong, Jia-Li Cao, Tian-Ying Zhang. Drafting of the manuscript: Tian-Ying Zhang, Jia-li Cao, Hua-Long Xiong. Critical revision of the manuscript for important intellectual content: Sheng-Xiang Ge, Jun Zhang, Quan Yuan, Tian-Ying Zhang, and Ning-Shao Xia. Technical or material support: Shao-Juan Wang, Wang-Heng Hou, Ya-Li Zhang, Bao-Rong Fu, Xiang-Yang Yao, Zhi-Yong Li and Ting Yang. Study supervision: Bao-Ying Huang, Zhi-Yong Li, Tian-Ying Zhang and Ning-Shao Xia.

## Patient consent for publication

not required.

## Ethics approval

The study was approved by the Medical ethics committee of School of Public Health of Xiamen University.

## Acknowledgements

We sincerely thanks for the team of Prof. Yong-Zhen Zhang, who publishing the genome sequence of SARS-CoV-2 (GenBank: MN908947) in the early stage of the epidemic. This work was supported by the National Natural Science Foundation of China (Major Program: 81993149041), Science and Technology Major Project of the Fujian Province (2020YZ014001), Xiamen Science and Technology Major Project (3502Z2020YJ02) and National Key Research and Development Program of China (2016YFD0500301).

## Reference

[1] Phelan A L, Katz R, Gostin L O. The novel coronavirus originating in wuhan, china: Challenges for global health governance. JAMA [J], 2020,

[2] Li Q, Guan X, Wu P, et al. Early transmission dynamics in wuhan, china, of novel coronavirus-infected pneumonia. N Engl J Med [J], 2020, 382(13): 1199–1207.

[3] Hui D S, E I A, Madani T A, et al. The continuing 2019-ncov epidemic threat of novel coronaviruses to global health - the latest 2019 novel coronavirus outbreak in wuhan, china. Int J Infect Dis [J], 2020, 91(264–266.

[4] Lu R, Zhao X, Li J, et al. Genomic characterisation and epidemiology of 2019 novel coronavirus: Implications for virus origins and receptor binding. Lancet [J], 2020, 395(10224): 565–574.

[5] Hofmann H, Hattermann K, Marzi A, et al. S protein of severe acute respiratory syndrome-associated coronavirus mediates entry into hepatoma cell lines and is targeted by neutralizing antibodies in infected patients. J Virol [J], 2004, 78(12): 6134–6142.

[6] Ou X, Liu Y, Lei X, et al. Characterization of spike glycoprotein of sars-cov-2 on virus entry and its immune cross-reactivity with sars-cov. Nat Commun [J], 2020, 11(1): 1620.

[7] Chen Y, Liu Q, Guo D. Emerging coronaviruses: Genome structure, replication, and pathogenesis. J Med Virol [J], 2020, 92(4): 418–423.

[8] Hoffmann M, Kleine-Weber H, Schroeder S, et al. Sars-cov-2 cell entry depends on ace2 and tmprss2 and is blocked by a clinically proven protease inhibitor. Cell [J], 2020,

[9] Yan R, Zhang Y, Li Y, et al. Structural basis for the recognition of sars-cov-2 by full-length human ace2. Science [J], 2020, 367(6485): 1444–1448.

[10] Ogino M, Ebihara H, Lee B H, et al. Use of vesicular stomatitis virus pseudotypes bearing hantaan or seoul virus envelope proteins in a rapid and safe neutralization test. Clin Diagn Lab Immunol [J], 2003, 10(1): 154–160.

[11] Fukushi S, Mizutani T, Saijo M, et al. Vesicular stomatitis virus pseudotyped with severe acute respiratory syndrome coronavirus spike protein. J Gen Virol [J], 2005, 86(Pt 8): 2269–2274.

[12] Nie J, Li Q, Wu J, et al. Establishment and validation of a pseudovirus neutralization assay for sars-cov-2. Emerg Microbes Infect [J], 2020, 9(1): 680–686.

[13] Letko M, Marzi A, Munster V. Functional assessment of cell entry and receptor usage for sars-cov-2 and other lineage b betacoronaviruses. Nature Microbiology [J], 2020, 5(4): 562–569.

[14] Schwegmann-Wessels C, Glende J, Ren X, et al. Comparison of vesicular stomatitis virus pseudotyped with the s proteins from a porcine and a human coronavirus. J Gen Virol [J], 2009, 90(Pt 7): 1724–1729.

[15] Whitt M A. Generation of vsv pseudotypes using recombinant deltag-vsv for studies on virus entry, identification of entry inhibitors, and immune responses to vaccines. J Virol Methods [J], 2010, 169(2): 365–374.

[16] Keyaerts E, Vijgen L, Maes P, et al. In vitro inhibition of severe acute respiratory syndrome coronavirus by chloroquine. Biochem Biophys Res Commun [J], 2004, 323(1): 264–268.

[17] Yang C, Compans R W. Analysis of the cell fusion activities of chimeric simian immunodeficiency virus-murine leukemia virus envelope proteins: Inhibitory effects of the r peptide. J Virol [J], 1996, 70(1): 248–254.

