## Supplementary file for "Robust neutralization assay based on SARS-CoV-2 S-bearing vesicular stomatitis virus (VSV) pseudovirus and ACE2-overexpressed BHK21 cells"

Supplementary figure 1：

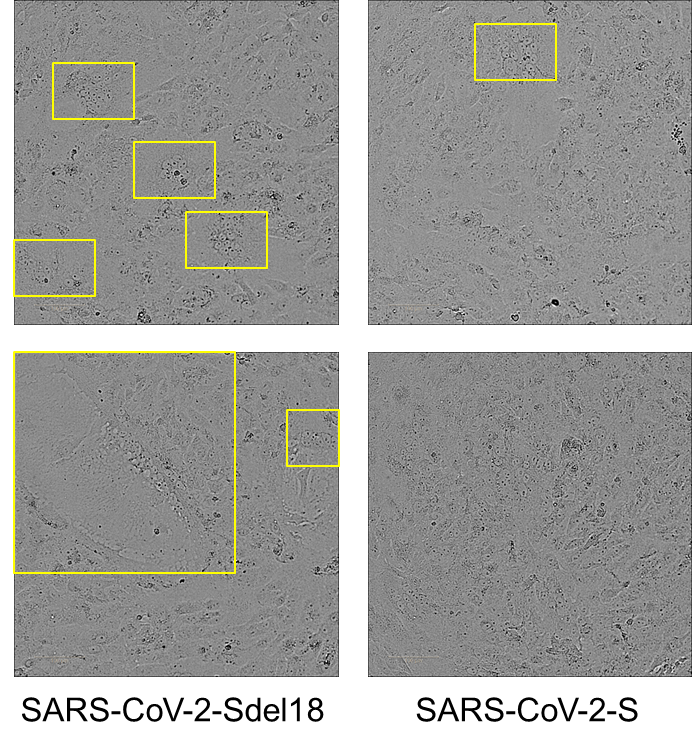

Supplementary figure 1: Formation of cell fusion produced by SARS-CoV-2-Sdel18 spike proteins compares to SARS-CoV-2-S. Vero-E6 was transfected with pCAG-nCoVS or pCAG-nCoVSde18 and observed the formation of symplasm 48 h post transfection.

Supplementary figure 2：

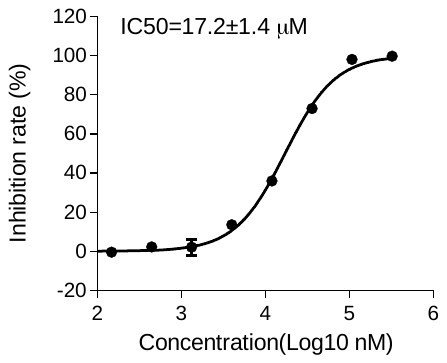

Supplementary figure 2: The dose-response of chloroquine in VSVdG-SARS-CoV-2-Sdel18 system. BHK21-hACE2 was incubated with VSVdG-SARS-CoV-2-Sdel18 virus (MOI=0.05) and different concentration of chloroquine. The fluorescence of cell was detected 12 h post infection. And the GFP positive cell number was counted to calculate the inhibit ratio. The IC50 was analysed by nonlinear regression.
